# A scale-free, fully connected global transition network underlies known microbiome diversity

**DOI:** 10.1101/2020.11.11.376103

**Authors:** Gongchao Jing, Yufeng Zhang, Lu Liu, Zengbin Wang, Zheng Sun, Rob Knight, Xiaoquan Su, Jian Xu

**Affiliations:** Single-Cell Center, CAS Key Laboratory of Biofuels and Shandong Key Laboratory of Energy Genetics, Qingdao Institute of BioEnergy and Bioprocess Technology, Chinese Academy of Sciences, Qingdao, Shandong, China; College of Computer Science and Technology, Qingdao University, Qingdao, Shandong, China; University of California, San Diego, CA, USA; University of Chinese Academy of Sciences, Beijing, China

## Abstract

Microbiomes are inherently linked by their structural similarity, yet the global features of such similarity are not clear. Here we propose as solution a search-based microbiome transition network. By traversing a composition-similarity based network of 177,022 microbiomes, we show that although the compositions are distinct by habitat, each microbiome is on-average only seven neighbors from any other microbiome on Earth, indicating the inherent homology of microbiome at the global scale. This network is scale-free, suggesting a high degree of stability and robustness in microbiome transition. By tracking the minimum spanning tree in this network, a global roadmap of microbiome dispersal was derived that tracks the potential paths of formulating and propagating microbiome diversity. Such search-based global microbiome networks, reconstructed within hours on just one computing node, provide a readily expanded reference for tracing the origin and evolution of existing or new microbiomes.

## Background

Microbiome composition, a fundamental feature of all microbiota in nature, is shaped by a plethora of environmental factors such as habitats, geographic locations, temperature, oxygen level and even day length [1, 2]. However, it remains unclear whether and how compositional changes at the “community to community” level among microbiomes are linked to the origin and evolution of global microbiome diversity [3-5]. For example, did microbiomes from different environments emerge and develop separately, or did the global microbiome start homologically and then spread to other habitats with compositional dispersal and dynamics? Over recent years, a large number of microbiome samples (e.g., Human Microbiome Project [6], Earth Microbiome Project [1], Tara Ocean [7], etc), mainly in the form of 16S rRNA amplicons, have been produced and accumulated [4]; however, the ability to cluster and model microbiomes at the global scale has however been hindered by the enormous volume and sheer complexity of such data (e.g. a distance matrix of 100,000 microbiomes contains ∼5*10^9^ elements).

## Results

### Microbiome transition model and search-based network

Here we describe the compositional dynamics and variation among microbial communities by a *microbiome transition* model. In this model, a microbial community is essentially a combination of microorganisms, and the structure of a community can be modified to another form by adding and/or removing species by compositional dispersal or fusion [8, 9] (**Fig. S1**). Theoretically, higher similarity between two communities indicates higher probability for such microbiome transition since fewer compositional exchanges are needed; however, it is not clear what level of similarity may indicate such microbial transition with reasonable confidence. Based on a pairwise full permutation of similarity calculation among all microbiome samples from the Microbiome Search Engine database (MSE [10]; 177,022 samples in total; refer to “**Microbiome sample collection**” for details) using Meta-Storms algorithm [11, 12] (**Table 1**), we consider that *direct transition* possibly exists between sample pairs with significant similarities that cause permutation *p*-value < 0.01 (*equation 1;* **Fig. 1**). As the result, we define the Meta-Storms similarity of 0.868 as the threshold for direct transition between microbiomes. By further analyzing the pairwise similarity in each habitat, we found that the threshold similarity of 0.868 is significantly high in the between-habitat similarity distribution (*p*-value=0.0022; **Fig 1B**); moreover it is higher than the upper boundary of most within-habitats similarities (17 of 20; **Fig 1C**). Thus the similarity threshold of 0.868 is sufficiently stringent for defining microbiome transition among the ecosystems.

**Table 1.**
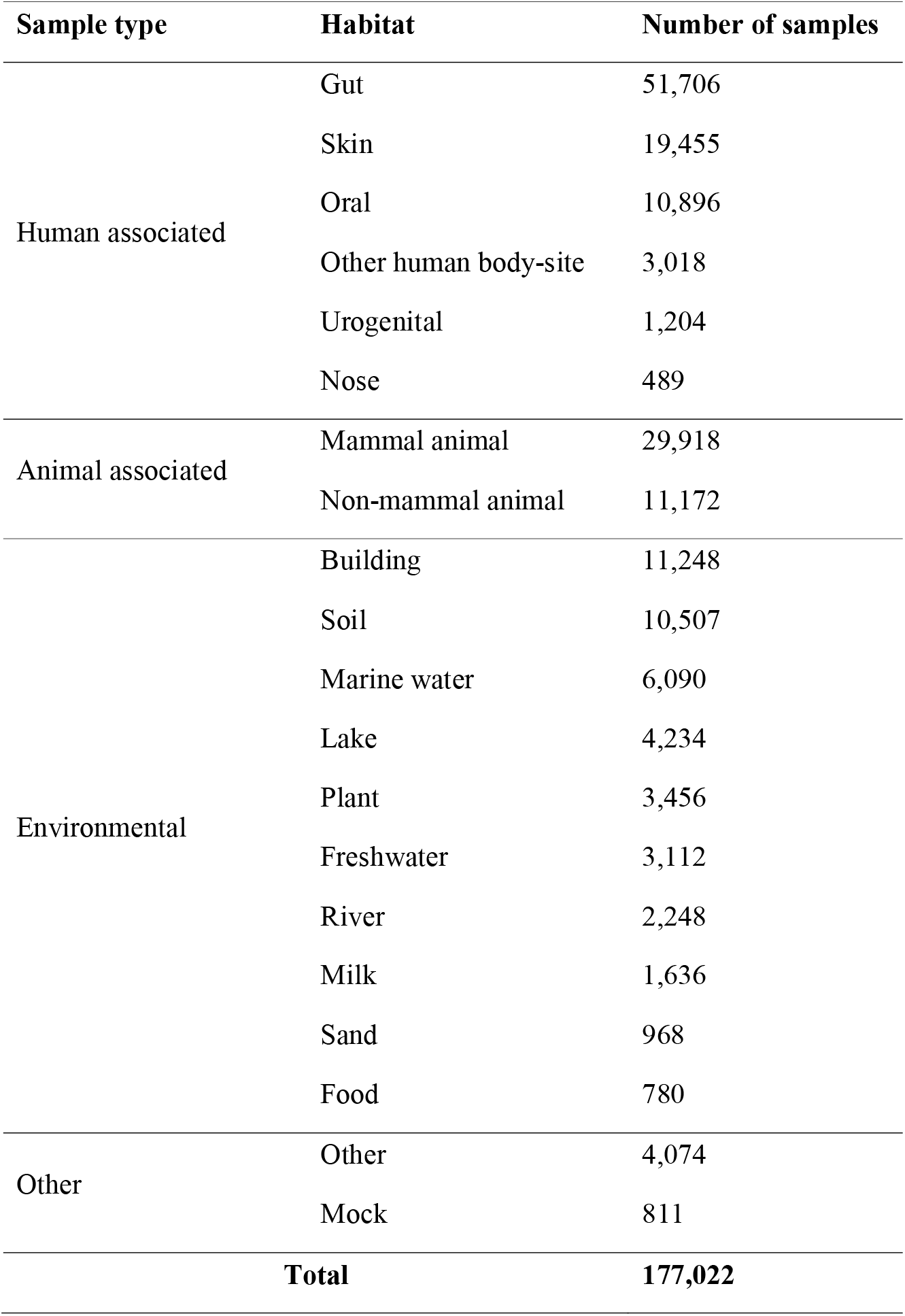
The distribution of samples among the habitats.

**Fig. 1.**
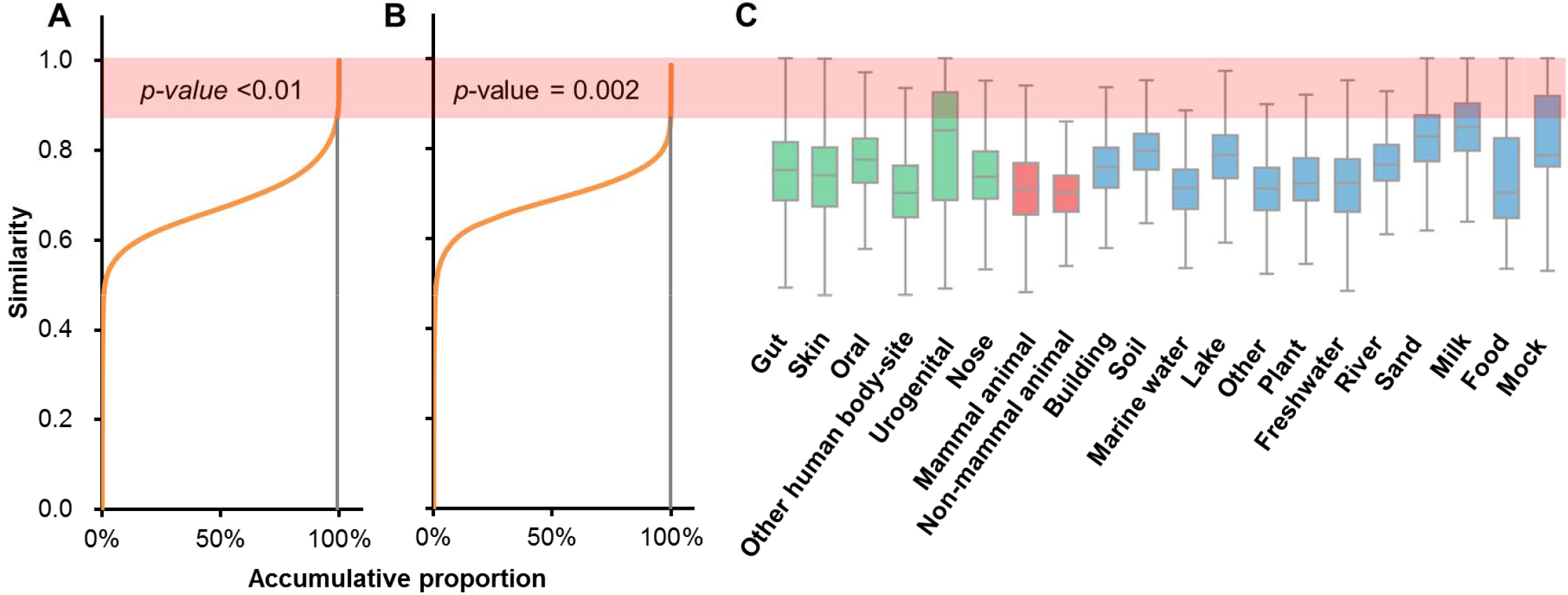
Distribution of pairwise similarity in *N*=177,022 microbiome samples. (**A**) *P*-value<0.01 of significant similarity values in the permutation determine the threshold of 0.868 (under the shadow) for putative direct transition. (**B**) The threshold has *p*-value=0.0022 among between-habitat similarity distribution. (**C**) The threshold is higher than the upper boundary of within-habitat similarities for most habitats. Three panels use the same Y-axis. *P*-values are calculated by permutation test.

Then for each of the input 177,022 microbiomes, we applied MSE to search against all other samples and find the top matches with similarity higher than 0.868. Based on the search results we constructed a transition network, in which each node is a microbiome, and each edge represents a direct transition (*equation 2*; **Fig. S2**). Collectively the network consists of 177,022 nodes (samples) and 11,175,742 edges (each called a *direct transition*). Notably, a pair of samples with low similarity can be connected via multiple edges (i.e., via a series of direct transitions across intermediate transfer samples), and such a sample pair is termed an *indirect transition* (*equation 3*).

### The transition network predicts microbiome habitat at a global scale

At the global scale, it is still not clear whether (and to what degree) similarity in microbiome structure implies similarity in ecosystem features [13, 14]. To quantitatively tackle this question, we compared the direct transition frequency between within-habitat (transitions of sample pairs from the same habitat) and between-habitat (transitions of sample pairs between two different habitats) cases in the transition network. For each habitat, the direct transition frequency is calculated by the average number of direct transitions per samples in this habitat. Notably, the direct transition exists more frequently between samples in the same habitat (**Fig. 2A**; two-tailed pair *t*-test *p*-value < 0.01). Thus the source environment of microbiomes dominates the microbial composition. We next used the transition network to predict the habitat (mock samples were not included) of each sample by its top neighbors (**Methods**). Via LOOCV (Leave-One-Out Cross Validation), 89.28% samples were correctly assigned by their original habitats (**Fig. 2B**; **Table 2**). Therefore, at a global scale, microbiome structure is strongly correlated with their environmental features.

**Table 2.** Prediction of habitat based on the microbiome network.

| **Habitat** | **Number of samples** | **Number of correctly predicted samples** | **Accuracy** | **Within-habitat transition frequency** | **Between-habitat transition frequency** |
| --- | --- | --- | --- | --- | --- |
| Gut | 51,706 | 50,431 | 97.53% | 66.89 | 5.40 |
| Skin | 19,455 | 17,464 | 89.77% | 52.11 | 17.82 |
| Oral | 10,896 | 10,070 | 92.42% | 55.10 | 10.98 |
| Other human body-site | 3,018 | 1,777 | 58.88% | 24.64 | 37.92 |
| Urogenital | 1,204 | 1,046 | 86.88% | 44.16 | 17.29 |
| Nose | 489 | 91 | 18.61% | 5.00 | 55.29 |
| Mammal animal | 29,918 | 28,010 | 93.62% | 50.67 | 8.87 |
| Non-mammal animal | 11,172 | 8,077 | 72.30% | 19.81 | 19.95 |
| Building | 11,248 | 8,942 | 79.50% | 40.24 | 24.54 |
| Soil | 10,507 | 9,978 | 94.97% | 54.49 | 7.28 |
| Marine water | 6,090 | 3,960 | 65.02% | 24.29 | 17.77 |
| Lake | 4,234 | 3,983 | 94.07% | 49.64 | 7.35 |
| Plant | 3,456 | 3,127 | 90.48% | 46.91 | 15.16 |
| Freshwater | 3,112 | 1,671 | 53.70% | 22.51 | 28.66 |
| River | 2,248 | 2,011 | 89.46% | 36.84 | 15.55 |
| Milk | 1,636 | 1,565 | 95.66% | 55.21 | 8.23 |
| Sand | 968 | 864 | 89.26% | 44.72 | 23.54 |
| Food | 780 | 677 | 86.79% | 39.01 | 26.57 |
| Other | 4,074 | 3,573 | 87.70% | 38.88 | 13.64 |

**Fig. 2.**
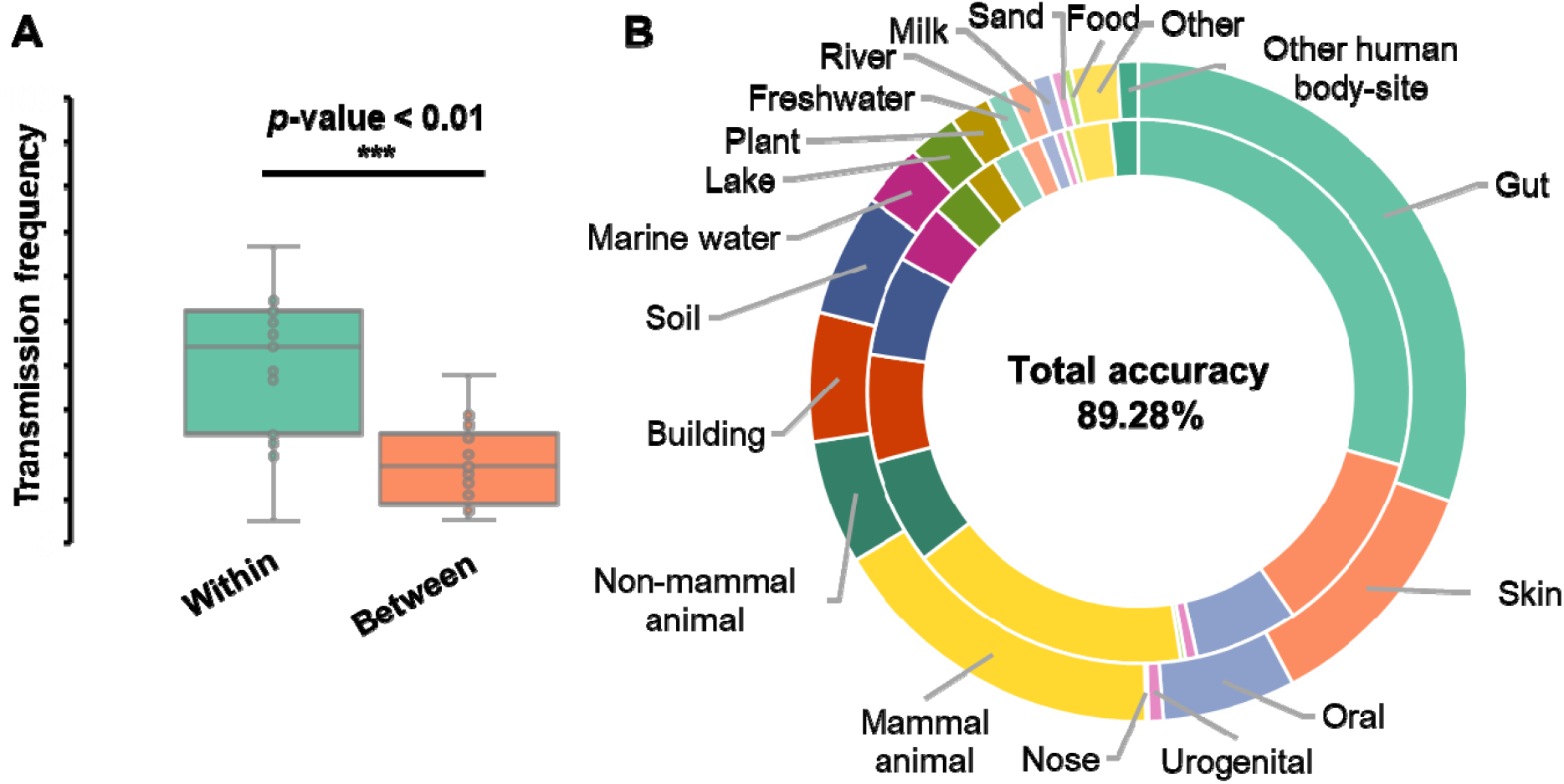
Global microbiome network predicts the microbiome habitat. (**A**) Frequency of within-habitat direct transition is significantly higher than that of between-habitat. *P*-value is calculated by two-sided *t*-test. (**B**) Habitat of 89.28% samples is correctly predicted by the microbiome network. Inner ring represents the proportion of real habitats and the outer ring is the proportion of predicted habitats.

“Mismatches”, i.e., microbiomes that were assigned to an incorrect habitat as predicted by the transition network, represent 10.72% (18,894 of 176,211) of all samples. Such mismatches are interesting as they can be caused by, and thus potentially indicate, frequent contact and interchanges of microbiota among the habitats. Those among human body sites are the most frequently observed mismatches (1.86% of all samples, same as bellow) caused by the daily contact and exchange of microbial composition [15]. Matches between non-mammal animal (sponge) and marine water are the second most frequent mismatches (1.76%). Mismatches across “Human Skin”, “Animal (pet)” and “Building (indoor environment)” represent 1.68% of samples, likely due to their sharing of the indoor environment (where the microbiome was largely sourced from humans [5, 16]). Moreover, 0.81% of the mismatches are between human-gut and mammal-animal-gut, which can be explained by the close phylogenetic relationship between human and other mammals and the coevolution of mammals and their gut microbiota [17]. Furthermore, mismatches are also observed (0.14%) where lake water samples are predicted as river water (note that this is the source stream of lake), or vise versa. Therefore, although the microbiome structure at the global scale is mainly shaped by their habitat, microbiome structure can be altered by, and thus reflect, the contact and exchange of microbiota from different environments.

### Microbiomes are connected globally by the transition network

The beta diversity of global microbiome may have evolved via two scenarios: (*i*) “polyphyly”, where microbiomes from different environments were generated and developed separately (**Fig. S3A**), or (*ii*) “monophyly”, in which microbiomes started homologically and then were dispersed to other habitats (e.g., via compositional transition, exchange or fusion; **Fig. S3B**). To distinguish between the two scenarios, we used the transitive closure algorithm to examine the connectivity of this transition network (**Methods**). A closure is a set of nodes (microbiomes), in which each microbiome can traverse to any other one by direct or indirect transitions (with finite steps). Hence being in a closure implies likelihood of transformation among samples via compositional exchange. Traversing all nodes in the network via the transitive closure algorithm revealed that 98.31% samples (174,032 of 177,022) can be clustered into a single closure (also named as the *main closure*). Under a condition that microbiota composition is distinct by habitat for 89.28% of all samples (**Fig. 2B**), such high connectivity suggests that the likelihood of polyphyly should be very low (probability < 1.5e-05; estimated by *equation 5*), and supports the monophyletic origin of global microbiomes and the formation of new microbiomes via such transitions (**Fig. S3B**). Notably, 1.69% (2,990 of 177,022) samples are still not included in the *main closure*, and they were mostly due to statistical inaccuracy (1.47% exhibit a similarity level that is only slightly below the threshold for being recruited into the *main closure*; *p-*value between 0.01 and 0.05) or curation errors (e.g. 0.16% are labeled as microbiome but actually pure-cultures or 18S/ITS amplicon samples). Therefore, the monophyly hypothesis best explains the origin and evolution of present-day microbiome structures.

To size the global microbiome network, we computed the pairwise shortest transition steps of all sample pairs in the *main closure* using the *Dijkstra* algorithm [18] (**Methods**). Interestingly, like the “small world” principle for social network [19], the microbiome transition network follows the “7-degree of separation” pattern (**Fig. S4A**). Specifically, any two microbiota in the *main closure*, even if they were sampled from different habitats and exhibit low similarity, can traverse from one to the other with only seven direct transitions on average [20], and 32 such steps at the maximum (i.e., the network diameter; **Fig. S4B**). Such a pattern underscores the high connectivity and thus surprisingly close interaction among microbiomes from diverse habitats at the planetary scale.

### The global microbiome network is scale-free and the connectivity is robust

Notably, in this global transition network, for each node, its edge degree *k* (number of direct transition neighbors) follows a Poisson distribution (**Fig. 3A**; Pearson *r*=-0.836 between *log(P(k))* and *log(k)*, i.e., *P*(*k*) ≈ *k*^−*γ*^), suggesting that the network is scale-free [21, 22]. One key feature of a scale-free network is the stability of topology, i.e., robustness to node removal from the network. To test the robustness, we removed different numbers of randomly selected samples and their associated edges, then assessed both size and leftover sample rate of the *main closure* in the residual network (percentage of residual nodes ranged from 5% to 95%; each such removal procedure was repeated for 10 times; **Methods**). When the number of nodes in the network reaches 80,000 (45% of total nodes), the connectivity rate curve of the *main closure* already exhibits a flat trend with 97.19% samples (98.31% prior to sample removal; **Fig. 3B**), and moreover, the mean transition steps and maximum transition steps (diameter) converge to 8 and 33, respectively (which were 7 and 32 before sample removal; **Fig. 3C**). Thus these parameters are quite stable, and not dependent on the increase of total sample number in the network. These findings suggest the robustness of microbiome diversity and similarity pattern among ecosystems at the global scale.

**Fig. 3.**
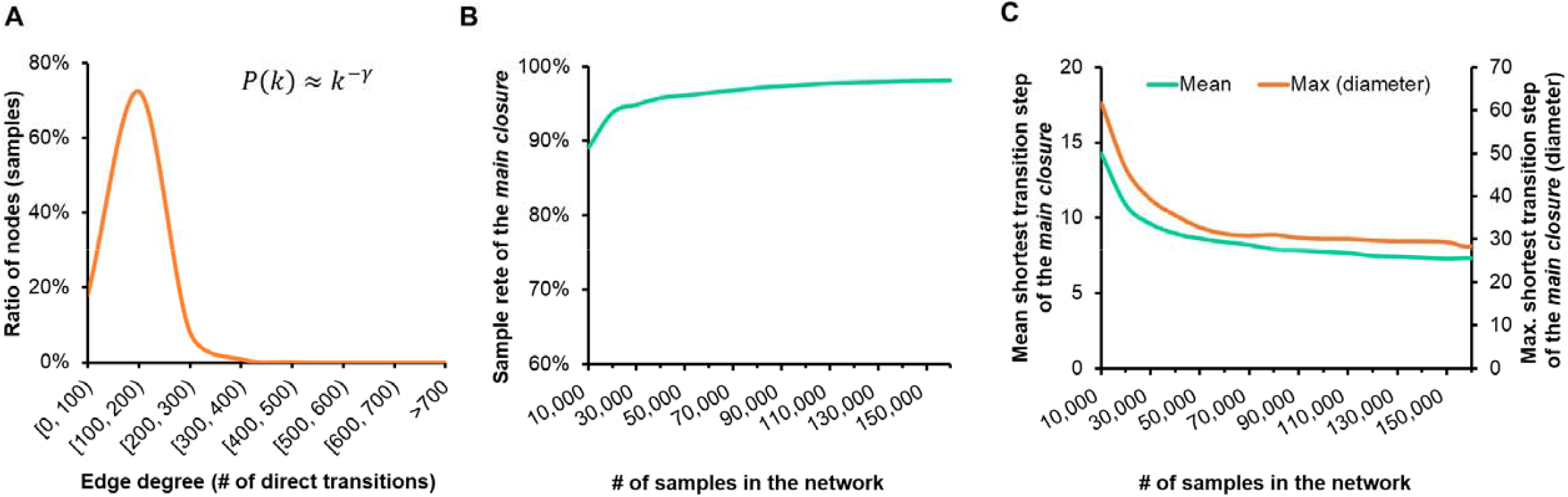
Robustness of the global microbiome network. **(A)** Node degree (number of linked neighbors) of the network follows the Poisson distribution, suggesting the network is scale-free. The effect of random node removal on the *main closure* in (**B**) sample rate, (**C**) mean shortest transition step and maximum transition step (diameter).

### The microbiome transition roadmap simulates the development of global microbial diversity among multiple ecosystems

As microbiome compositions are dominantly determined by their habitats, the full connectivity of global microbiomes in the network suggests the ability to reconstruct how the microbial diversity spread among different habitats at a macroscopic scale. This “microbial dispersal” roadmap can be simulated by a sub-network that *i)* covers and links all samples, and *ii)* consists of deterministic finite transition steps without cyclic or redundant routes. Thus we derived such a roadmap (**Fig. 4A**) by parsing the Minimum Spanning Tree (MST) of the *main closure* using the *Kurskal* algorithm [23] (**Methods**). As the global optimum with the highest overall transition probability (similarity), the MST maximally captures the transition pattern of world-wide microbial diversity among all the 19 habitats (with the “mock” samples excluded). For example, marine microbiomes most probably exchange with two other environments: one is sand which is geographically close to the shore, while the other is non-mammal animals such as fishes. These observations also suggest that sand and freshwater microbiomes are the “gateways” to soil, plants and human-associated habitats such as gut, oral, skin, and the human living environments.

**Fig. 4.**
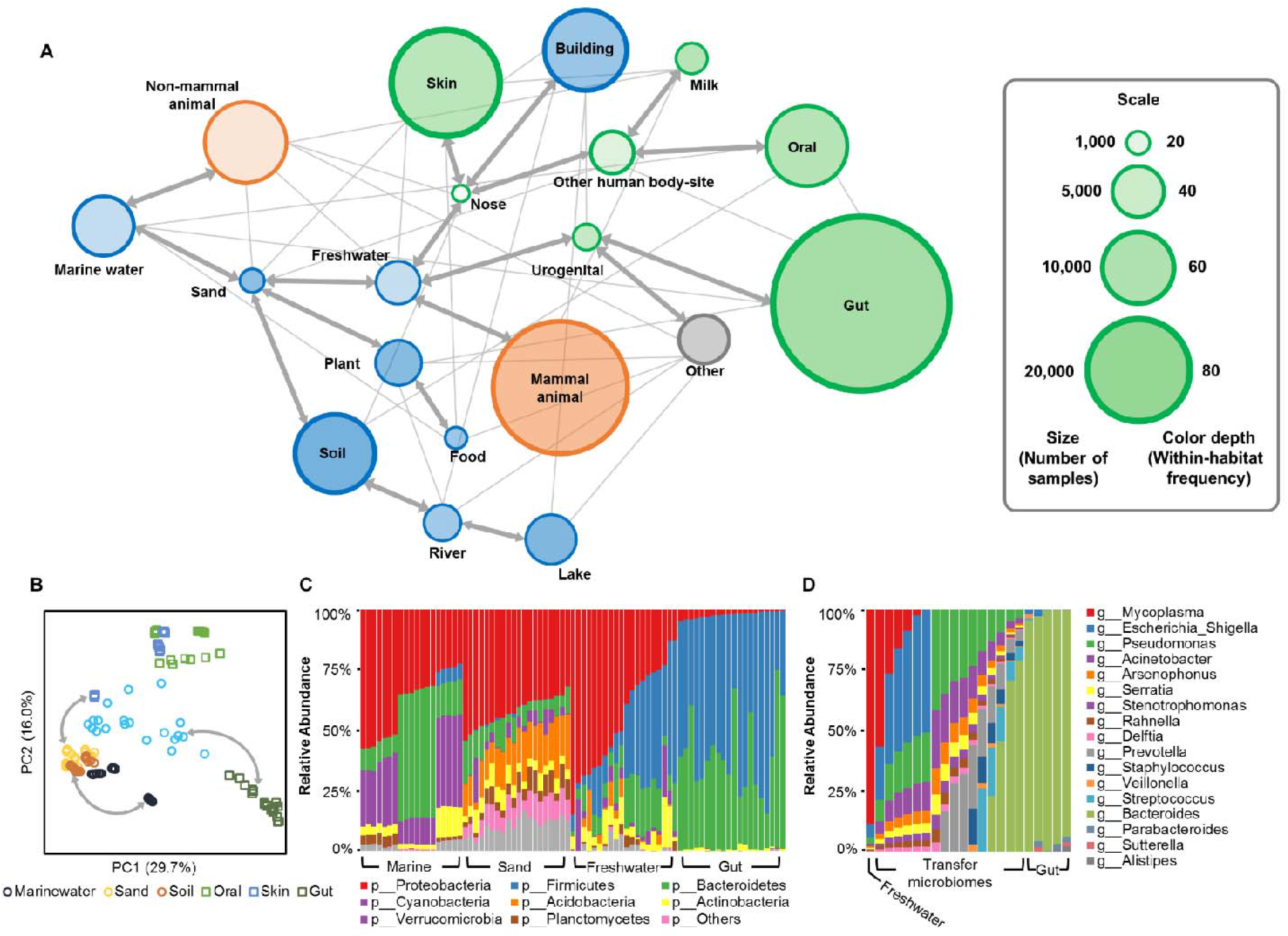
Roadmap of the global microbiome transition among habitats. (**A**) Bold arrow lines are the roadmap that represents the maximum overall similarity. The number of samples in each habitat is scaled by the node size, and the within-habitat transition frequency is represented by the node color depth (compared to the rim). (**B**) The PCoA parsed from a subset of 140 microbiomes demonstrates the roadmap by the equivalent topology. (**C**) The phylum-level compositional shift of a microbiome transition route for marine to gut environment. (**D**) The genus-level compositional shift of a transition case from a freshwater microbiome to gut samples.

This roadmap is verified by the isomorphic pattern of PCoA derived from a subset of 140 samples randomly selected from six habitats (**Fig. 4B**). Moreover, we employed a marine-gut route, which represents one of the longest transitions in this subset, to illustrate the high-resolution transition procedures (based on phylum-level compositional variations; urogenital and gut microbiomes were combined as they are very close in the global scale; **Fig. 4C**). Zooming in of this marine-gut route revealed a series of structure shifts that transform a freshwater microbiome to gut samples. Starting from an actual freshwater sample [24], in each step, organisms enriched in freshwater [25] (e.g. *Mycoplasma* and *Escherichia*) were removed/reduced, and organisms abundant in gut [26] (e.g. *Bacteroides* and *Parabacteroides*) were added/increased. Although a single step might have caused just slight modification on the microbiome structure, after several iterations this sample can be smoothly transited to gut microbiomes [27] in the network, via a series of transfer samples (**Fig. 4D**).

### Microbiome transition over time and across geography

To test the feasibility of modeling microbial dynamics by the global microbiome transition network, we used a longitudinal cohort to describe the transition of human microbiomes across time. In this dataset, 1,963 samples were collected from three body sites (gut, oral cavity and skin) of two individuals (I, male; II, female) over 396 time points [28]. Our search-based network analysis revealed that the microbiome composition of each body site exhibits significant variations across time (**Fig. 5A, B** and **C**; **Fig. S5**), while skin and oral microbiomes were clustered into the same closure by direct transition (**Fig. 5D, E and F**). These suggest that microbiome transition is ongoing within each site and between the skin and oral sites across different time points. In addition, for both hosts, gut samples are “isolated” from the skin-oral closure, consistent with the global microbiome transition map (**Fig. 4A**) where gut microbiomes are in a distinct route from skin and oral ones. Therefore, although oral cavity and gut are both of the digestive tract and moreover microbial translocation from oral to gut can occur [29], the oral microbiota have more likely been derived from or more prominently shaped by the skin microbiota (and vice versa) than from the gut microbiota. This seemingly counter-intuitive finding can actually lend support from the sharing of a more aerobic and less acidic environment by skin and oral cavity (pH and oxygen level are known to have large effect size in microbiome structure [1].

**Fig. 5.**
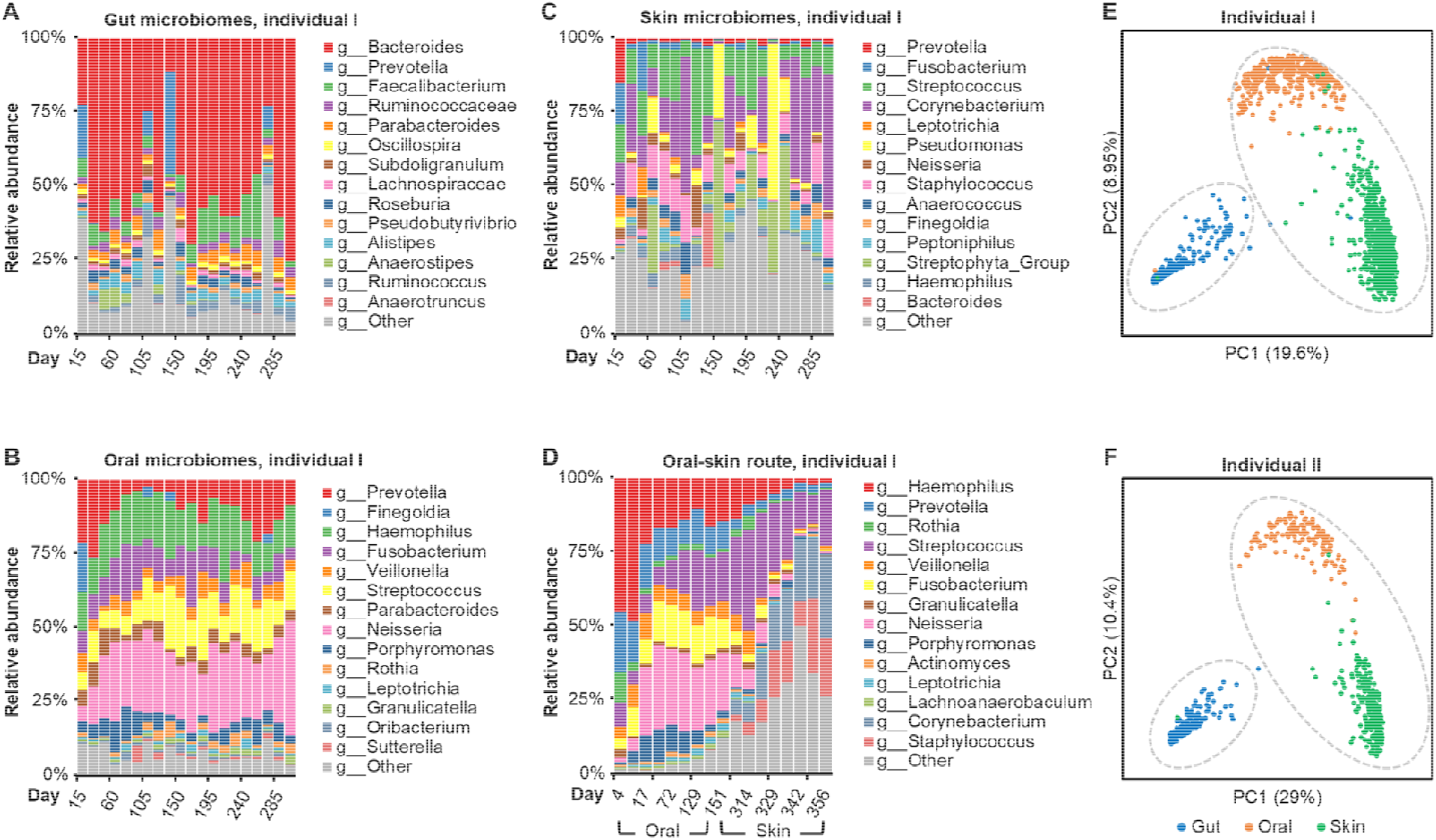
Transition of human microbiome across time and body sites. The within-habitat microbiome transition of gut (**A**), oral (**B**) and skin (**C**) of Individual I across 396 time points. (**D**) The oral-skin microbiome transition of individual I across time. Only selected samples were shown. The transition patterns of Individual II are shown in **Fig. S5. E** and **F**, PCoA of the two individuals’ time-series microbiomes: skin and oral microbiomes are linked in a closure by direct transition (highlighted by grey dotted line) and gut samples forming another closure.

On the other hand, to validate the connectivity of microbiomes from varied geographical locations, we constructed a search-based network by a single dataset [30] that contains 3,850 samples collected from six habitats (human gut, human oral, non-mammal animal, plant, soil and freshwater) and locations at North America (Urbana IL, Columbia MO, Aurora CO, Ithaca NY and Lansing MI, etc). With this dataset, the authors concluded that there was no overlap of abundant bacterial taxa between the microbial communities from human gut and plant roots [30]. Consistent with this conclusion, the network-based analysis based solely on this dataset, that samples were distributed into three isolated closures of direct transitions (**Fig. 6A**). However, once extra 1,635 samples from MSE database that connected the different closures of this local network were added, a single closure that covers 97.74% samples and integrates the original three closures emerged, with the newly included samples serving as “transfer nodes” (i.e., samples that link two clusters in network; *equation 3* in **Methods**) that provide additional indirect transitions(**Fig. 6B**; **Methods**). Notably, among such “transfer” microbiomes, most (96.89%) were from the same habitats as the original dataset, and others were mainly from sand and marine, which are found as the transfer nodes among non-mammal animal, plant and soil microbiomes in our global microbiome transition roadmap (**Fig. 4**). This example demonstrates that although microbiomes from diverse environments and isolated geographical locations can have very distinct structures, they can still be linked within a single closure in the microbiome transition network, i.e., evolve from each other, as long as global beta-diversity is adequately surveyed and covered. These results, which directly challenge the conclusion that multiple host microbiota compositions were independently evolved [30], underscores the importance of deriving or validating “local” datasets under the context of our global microbiome network, particularly when discussing similarity (i.e., beta-diversity), interaction or other kinds of relationship among microbiomes.

**Fig. 6.**
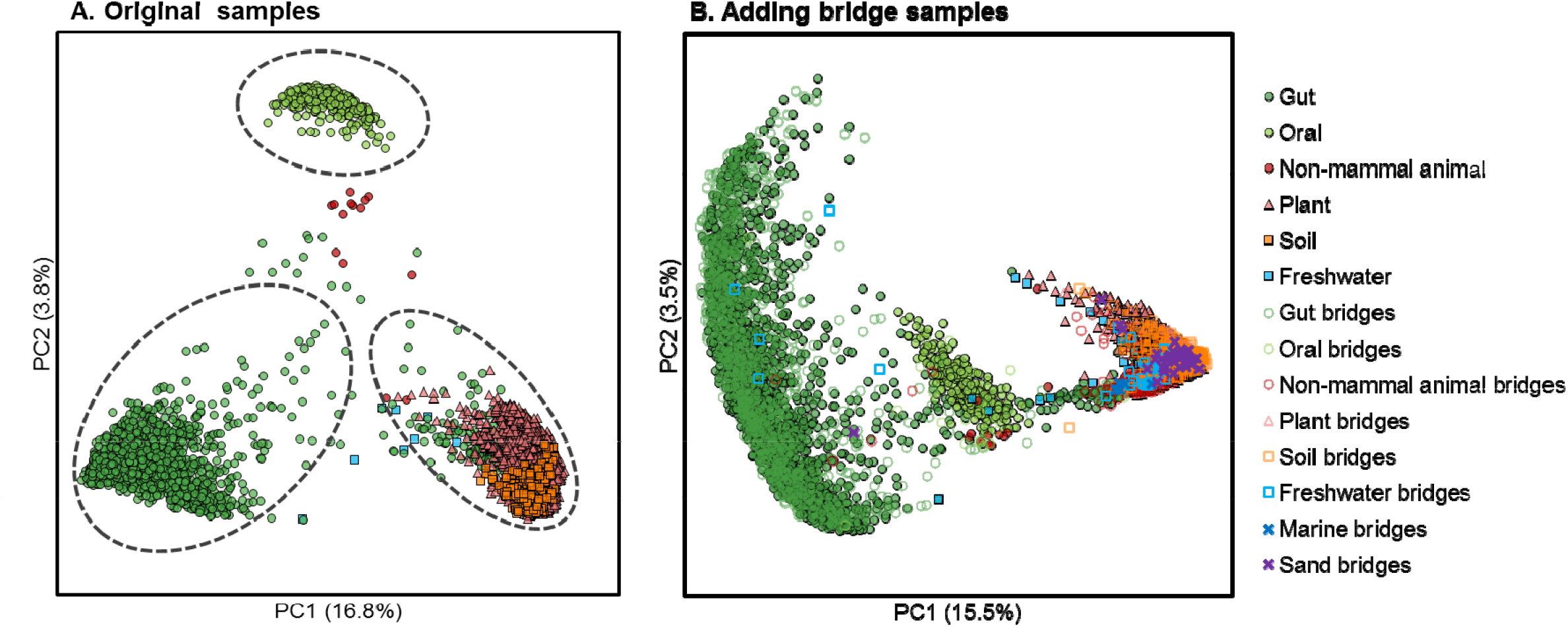
Microbiome transition across habitats and geographical locations. (**A**) The 3,850 samples from six habitats are included in three isolated transition closures, of which the sample proportions are 43.22%, 7.90% and 45.53%, respectively. (**B**) After adding extra 1,635 bridge samples from the MSE database, three closures are merged into a single closure by direct transition, which contains 97.74% samples.

## Conclusions and Discussions

Microbiota have been co-evolving with and shaping our planet, but their origin and evolution at the global scale remain elusive, due to the lack of fossils and the methodological challenges associated with integrating and mining such large-scale high-complexity data [14]. Although “species-to-species” interactions have been mapped by co-occurrence analysis on microbiomes across various habitats [31, 32], efforts to globally traverse and interrogate the vast microbiome data space at the “community to community” level have just started [10, 33, 34]. Here we propose a microbiome transition model and a network-based analysis framework to describe and simulate the variation and dispersal of the global microbial beta-diversity across multiple habitats. Benefited from the extremely-high search speed of Microbiome Search Engine [10], we introduced a global microbiome network with 177,022 microbiome samples that contains 11.3 billion sequences. By traversing such network, we showed that the microbiome structures are world-widely connected by significant similarity that follows the “small world” principle. This endeavor reveals the inherent homology of the global microbiome diversity and supports the monophyletic origin of all microbiomes on Earth. Then we drew the first global microbiome transition roadmap to illustrate the potential yet most likely paths that can explain the evolution process of global microbiomes.

Due to the ongoing exponential growth of microbiome sequencing data, current beta-diversity analysis approaches, which mainly rely on the *O(n*^*2*^*)-*complexity pairwise relations (*n* is the number of samples) such as Principle-Coordinate-Analysis (PCoA) and clustering, have become increasingly stressed or even impractical, particularly when computational resources are limited. Here we tackled this challenge via a search-based network, which is built on the “neighbors”, i.e., those with the highest similarity for each sample; this strategy reduces the computational complexity to *O(c*n)* (c is constant, i.e., the number of neighbors), and thus enables deciphering the pairwise similarity for > 100,000 microbiomes within 3 hours on a single computing node. As a result, the global microbiome transition roadmap, which will be regularly updated as community resource, can serve as a reference for interpreting or validating those existing or future observations on inter-microbiome similarity, association or interaction, since local microbiome datasets can be readily aligned to this global roadmap based on their shared nodes. Moreover, such a network-based analysis framework, which can be extended to shotgun metagenome datasets, provides a new perspective for tracking back or predicting microbiome evolution with fine resolution yet at the global scale.

## Methods and Materials

### Microbiome samples collection

We used all the microbiome samples from Microbiome Search Engine database (http://mse.ac.cn). Samples were collected from 572 studies/projects that including 20 habitats (**Table S1**). OTUs were picked and annotated against Greengenes [35] full length 16S rRNA gene sequences (version 13-8) on 97% similarity level by Parallel-META 3 [36] (version 3.4.4). Variation of 16S rRNA gene copy number was normalized based on the IMG/M database [37]. We set a minimum sequence number of 500 and minimum 16S rRNA mapping rate of 80% for each sample to ensure high quality of the reference datasets. Finally, *N*=177,022 samples with 11,302,841,991 mapped sequences assed the quality control and curation (**Table S1**).

### Calculation of pairwise microbiome similarity matrix for definition of direction transition

The pairwise similarity matrix of all *N*=177,022 samples was entirely permuted (totally (N * N-1)/2 = 15,668,305,731 times) to examine the distribution of microbiome phylogeny similarity using Meta-Storms algorithm [11, 12] in Parallel-META 3 software package. By setting a cutoff *p*-value < 0.01 in the permutation of the similarity (rank of top 1%), we got the Meta-Storms similarity 0.868 as the statistical threshold of the significant high value to define the direct transition (this threshold is also referred to as *T*_*d*.*t*_). Thus the transition model can be described in the following form

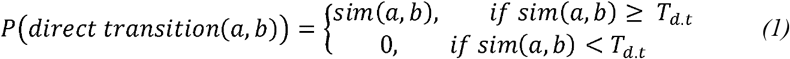

in which *s*_*i*_ and *s*_*j*_ are two arbitrary microbiomes, and *sim(s*_*i*_, *s*_*j*_) represents their Meta-Storms similarity.

### Search-based microbiome network

The search-based microbiome network is built using Microbiome Search Engine (MSE) [10]. For each sample we searched it against all other samples for the top 100 matches, and connected it with the matched samples that have similarity higher than the threshold of direct transition (*T*_*d*.*t*_=0.868). By iterating such search with all samples, we constructed a global network *G* (*equation 1*).

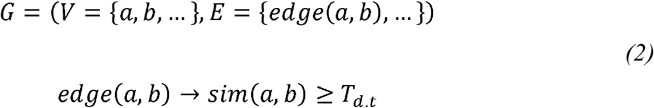

In this network, one node (e.g. *a* or *b* in *equation 2*) is a single microbiome, and edges (e.g. *edge(a, b)* in *equation 2*) that link the nodes are direct transitions (**Fig. S2**). Finally in the network there were 177,022 nodes (samples) and totally 11,175,742 edges (direct transitions). In this network, a pair of samples with low similarity can be connected by a path of multiple edges, i.e., an *indirect transition*:

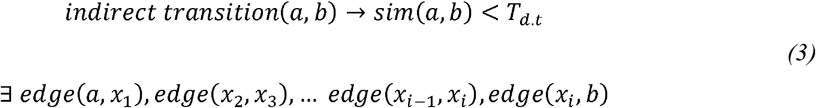

Then here *x*_*1*_, *x*_2_, …, *x*_*i*_ are defined as the “transfer samples” for that underlie the indirect transition from *a* to *b*.

### Prediction of habitat using microbiome network

In the network *G* we predicted the source habitat of each microbiome by its top *n*=10 neighbor samples and similarities. For an arbitrary microbiome sample *a* in the network, similarities to its top 10 neighbors are *S* = {*s*_1_,*s*_2_,…,*s*_*n*_}, while the *n* neighbor were from *m* (1 *≤ m ≤ n*) different habitats as *H* = {*h*_1_,*h*_2_,…,*h*_*m*_}, then the probability for the predicted habitat of microbiome *a* as *h*_*k*_ is calculated by

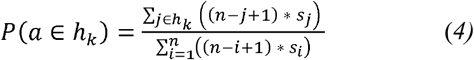

Here *j∈h*_*k*_ means the habitat of neighbor *j* is *h*_*k*_ (1 ≤ *k* ≤ m). Then the predicted habitat that has the highest probability *P* will be assigned to the sample *a* as the prediction results.

### Probability of global transition among all habitats in the network

By *equation 4* we calculated that at the global scale the microbiota composition is distinct by habitat, that the probability of transition among the same habitat is 89.28% (**Fig. 2B**). To calculate the overall probability of connecting all habitats in the transition network, we can start from connecting arbitrary two habitats of which the probability *p*_*transition*_*(n=2)* = 1-89.28%. When one more habitat added into the network, the probability of transitions among the three habitats can be calculated as *p*_*transition*_*(n=3)* = *p*_*transition*_*(n=2) \** (1-89.28%^2^), where the square of 89.28% represents the probability that no direct transition between the added habitat and the former two habitats. Then we can expand such procedure to estimate the probability for connecting *n* habitats in the transition network by

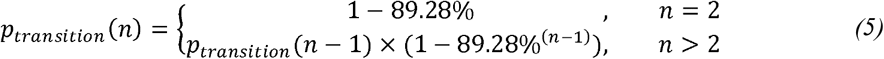

### Transitive closure algorithm of microbiome network

In the microbiome transition network, a closure is a subset of nodes (microbiome) are fully connected, that each microbiome is possible to link to any other sample by direct or indirect transitions (with finite transfer nodes). Closures can be initialized by an arbitrary node in the network, and then expanded by adding more external nodes that are directly connected with this closure (**Fig. S6**). If two or more closures are connected by any edge, these closures can also be merged as one closure. By the traversal among all nodes in the network *G* we get a *main closure C* with 98.31% samples.

### Size of the microbiome network

In the *main closure C*, there are always multiple routes between two indirectly connected nodes (microbiomes). We count the edge between two directly linked nodes as 1, so the length of indirect routes is the number of transfer nodes + 1 on this route (**Fig. S4**). We used the *Dijkstra* algorithm [18] by Python package *igraph* (0.7.1 running inside Python 3.6.1) to find the pairwise shortest transition steps (with smallest number of transfer nodes) between all indirectly linked node pairs in the *main closure C*. Thus the number of maximal steps among the shortest route is the diameter of the closure. The diameter means in this closure, any two microbiomes could be linked to each other by a route with steps that smaller than the diameter.

### Minimum spanning tree for the roadmap of microbiome network

In a transitive closure, a spanning tree is a sub-network that connects all nodes (microbiomes) with no cycle. For two directly linked samples *a* and *b*, we define their distance as

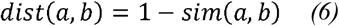

the Minimum Spanning Tree (MST) could be consider as the global transition path of all samples with the highest overall transition probability since it links all samples with the shortest total distance. In the *main closure C* we used the Kruskal algorithm [23] to calculate the 2-level MST to reflect the transition among different habitats from the global scale.

The first level MST was on “sample resolution”, based on which we then make the second MST on “habitat resolution”. Initially we calculated the first level MST of the *main closure C*, and then generated the habitat-based network *G’* (*equation 2*), each node represents one habitat, and distance between two habitats *h*_*i*_ and *h*_*j*_ is the average distance of all edges that linked the two habitats in the MST. Then we computed the second-level MST (*G’*), which illustrates the global microbiome transition roadmap across multiple habitats.

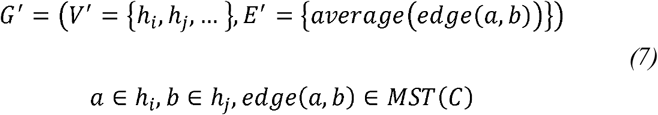

The significance of the roadmap (MST(*G’*)) was assessed by the permutation test of the topologically equivalent sub-network in the *main closure C* of the original network. Specifically, in a permutation, for each edge that connects two habitats (eg. *habitat*_*i*_ and *habitat*_*j*_) in the roadmap, we also randomly selected an edge that connects two samples (eg. sample *a* and *b*) respectively from these two habitats (eg. *a*∈*habitati* and *b*∈*habitat*_*j*_). Since we iterated the permutation for 10,000 times, if the total distance of the roadmap is smaller than 99% of permutated network (also means the total probability is in top 1%, *p*-value < 0.01), we can consider the roadmap MST(G’) is significant in the *main closure C*.

### Search-based sample selection from reference database to link separated closures

To select transfer samples from a reference database to link two separated closures, we search all samples of each closure against the referenced repository for top matches with higher similarity than the direct transition cutoff (*T*_*d*.*t*_=0.868), and the overlapped matches between the two closures are the transfer microbiomes that link the two closures. If there is no overlap in the matches, we then extend each of the closure by adding their matches, and repeat the search process until we find any transfer sample. On the other hand, once closures cannot be furtherly extended by database search but still no available transfer sample is found, this means that no sample in the reference database is able to work as the transfer node to link the two separated closures by direct transition.

## Acknowledgements

J.X. acknowledges support of grants 31327001, 31425002 and 81430011 from National Natural Science Foundation of China (NSFC), KFZD-SW-219-4 and ZDBS-SSW-DQC002-03 from Chinese Academy of Sciences (CAS). X.S. acknowledges support of grants 31771463 and 32070086 from NSFC. R.K. acknowledges support from the U.S. National Institutes of Health, the National Science Foundation, and the Alfred P. Sloan Foundation. We thank the Operations Research Community-QIBEBT for the inspiring discussions.

## Author contributions

J.X. and X.S. conceived the idea. X.S., and G.J., developed the software and algorithm. G.J., and Y.Z. performed analysis. L.L., Z.X., Z.W. and Z.S. contributed to data collection and curation. X.S and J.X. wrote the manuscript.

## Cod and data availability

All the source code and source data used in this work are available at https://github.com/qibebt-bioinfo/microbiomenetwork.

