## Supplemental figures for "A scale-free, fully connected global transition network underlies known microbiome diversity" (from another version of this preprint: version 1, 2020-11-11)

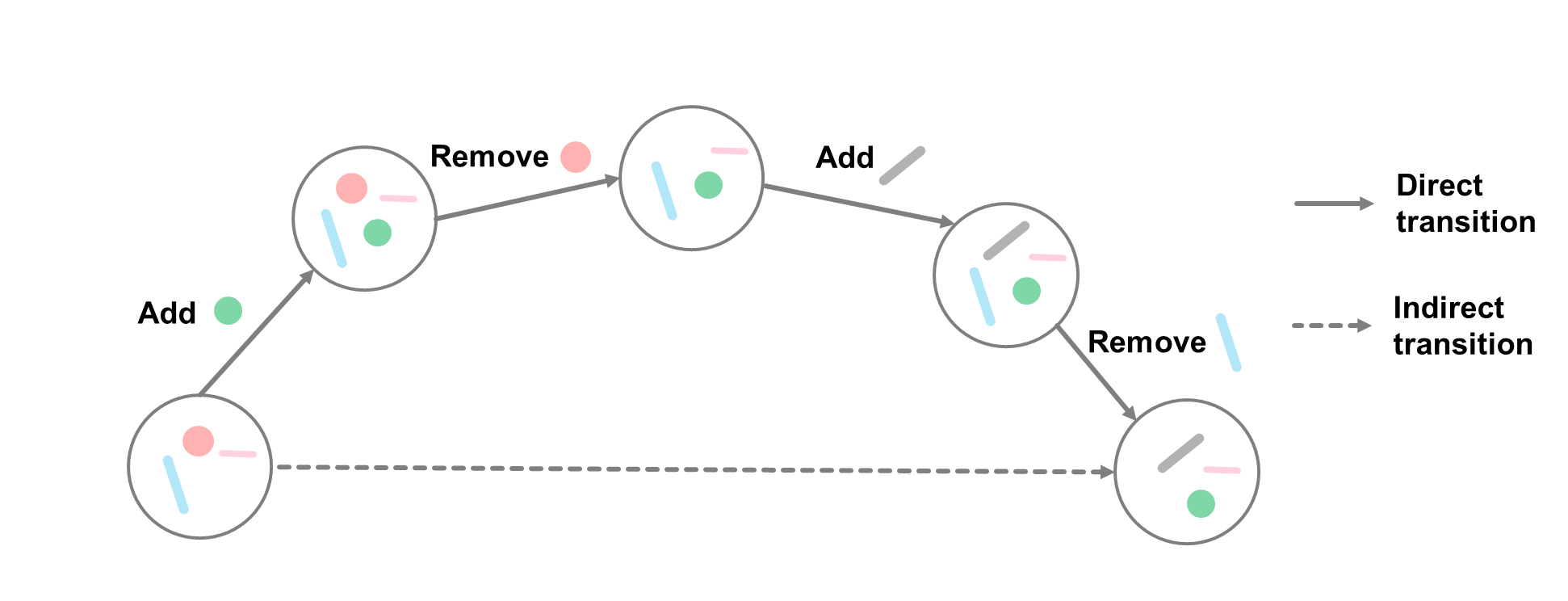

**Fig. S1. Microbiome transition model.** A microbial community is a combination of microorganisms, and the composition of one community could be modified to another form by increasing and/or decreasing the amount of its species, or adding and/or removing species from the community.

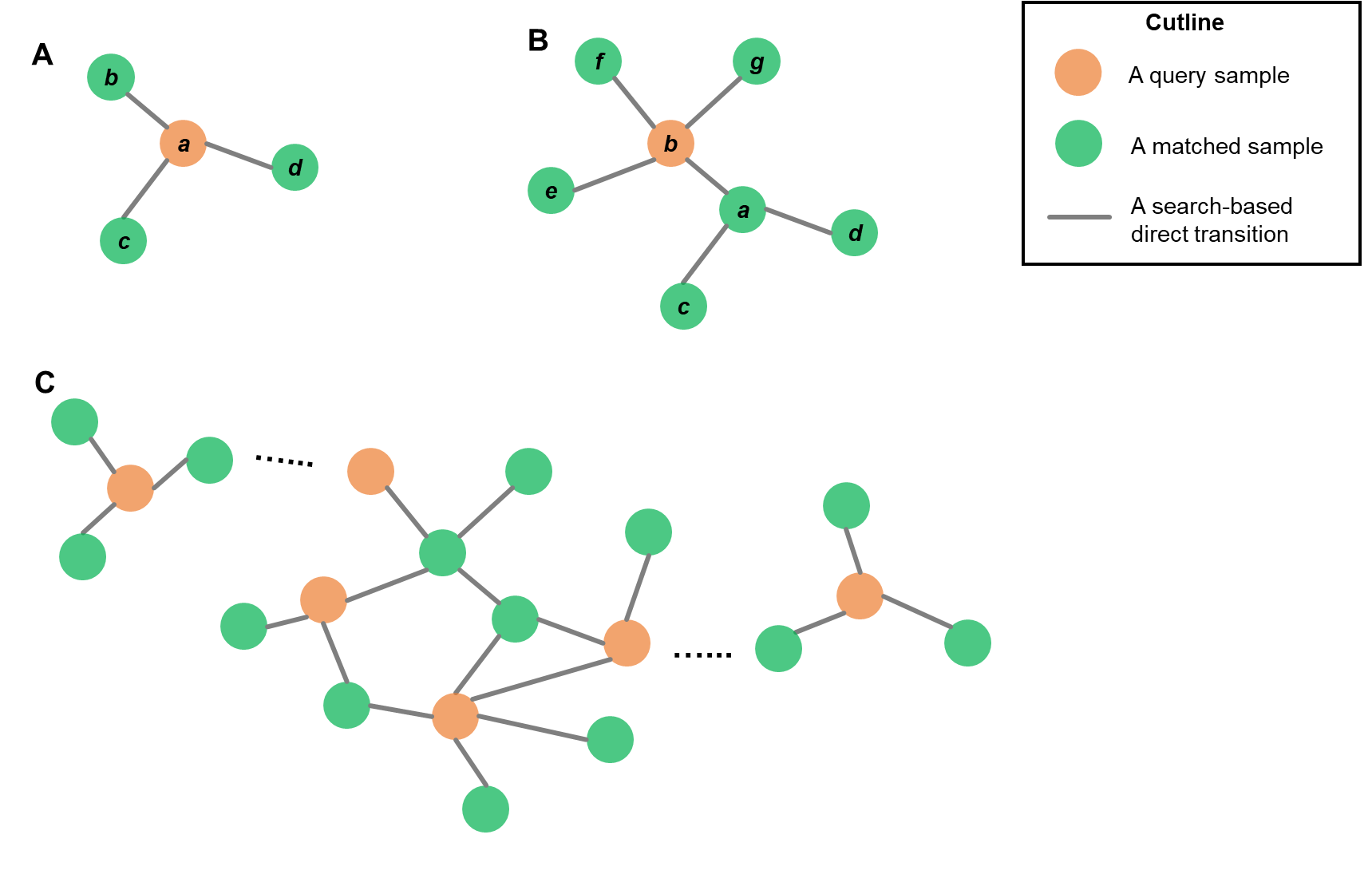

**Fig. S2. Construction of search-based microbiome transition network.** (**A**) For a query sample *a* we search it against all other samples, and connect it with the matched samples *b*, *c* and *d* that have similarity higher than the threshold of direct transition. (**B**) Then iteration this procedure to sample *b* and connect it with *e*, *f* and *g*. (**C**) By iterating the search through all samples we construct a global transition network.

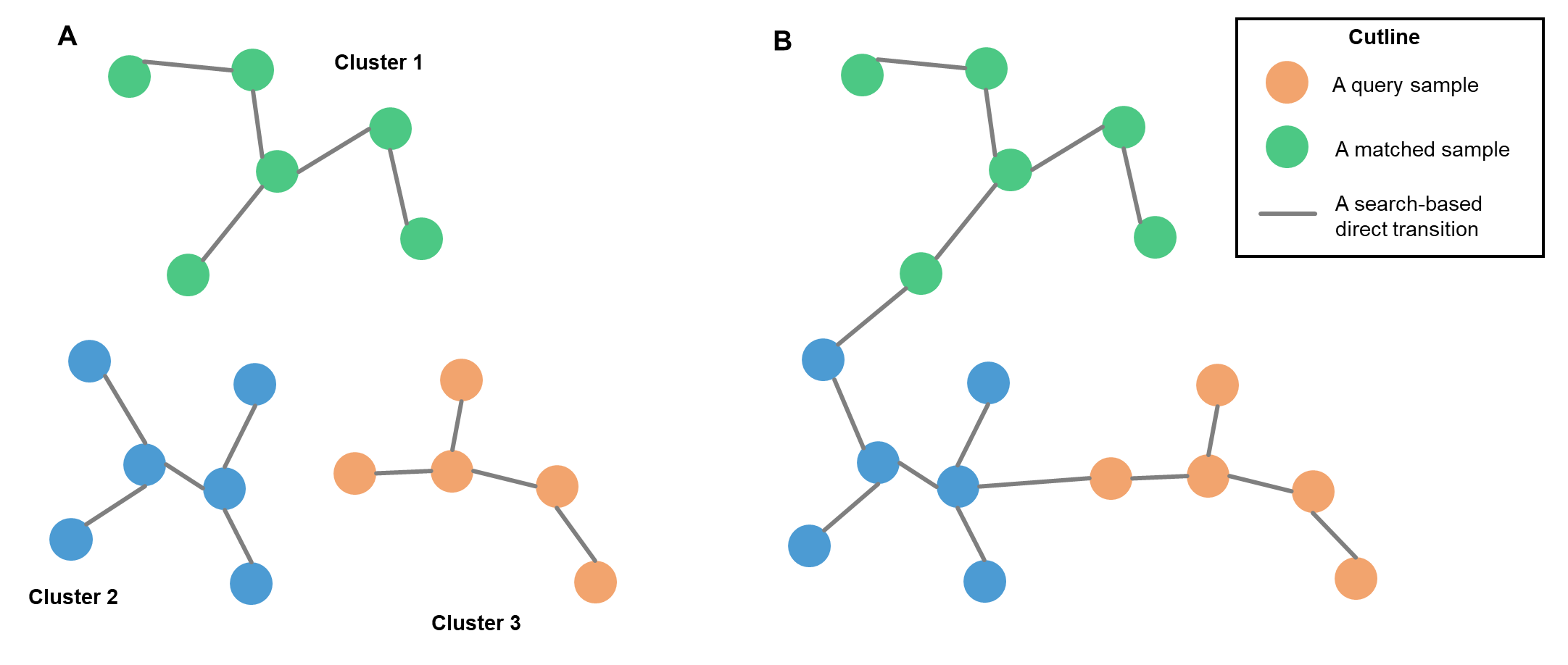

**Fig. S3. Two scenarios of the global microbiome generation.** (**A**) Microbiomes of different environments are generated and developed separately. (**B**) The global microbiome started from the same ancestors and then spread to other habitats with compositional transition and exchange.

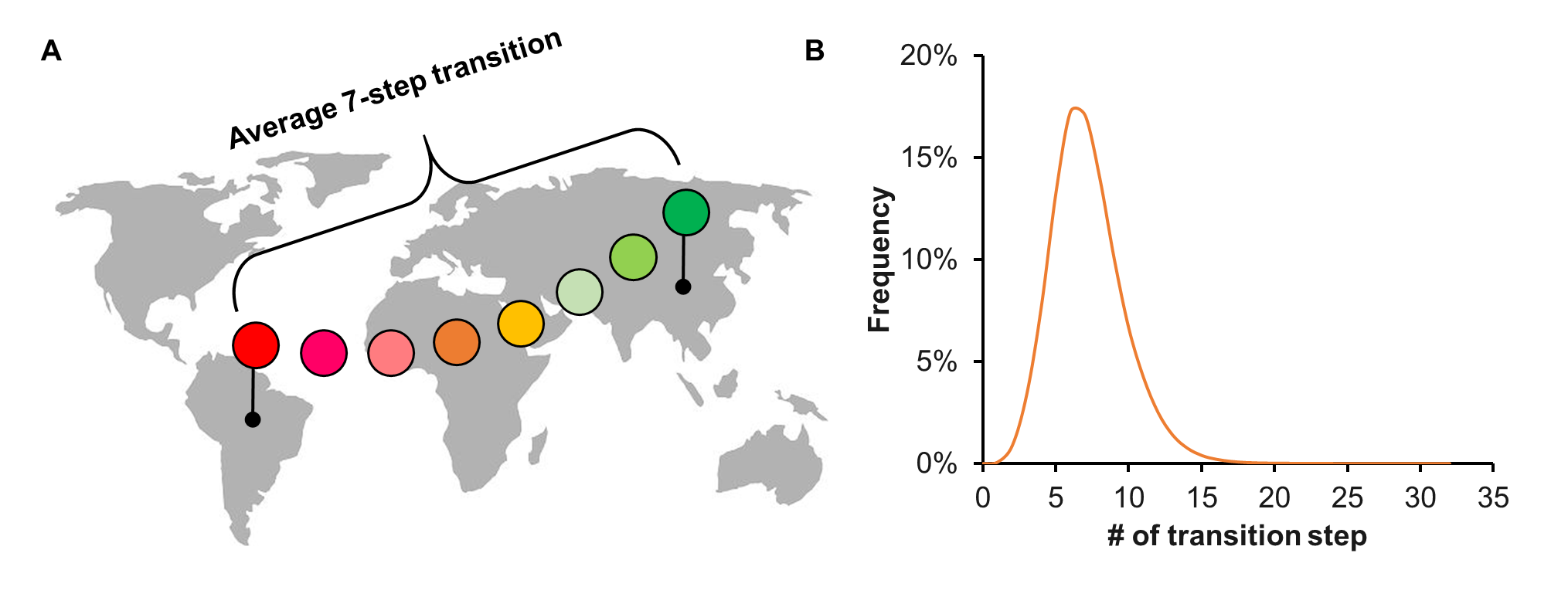

**Fig. S4. The “small world” principle of the microbiome transition network. (A)** Any two microbiomes in the *main closure* could be transited to each other by a 7-step transition on average. **(B)** Distribution of the pairwise shortest transition steps of all sample pairs.

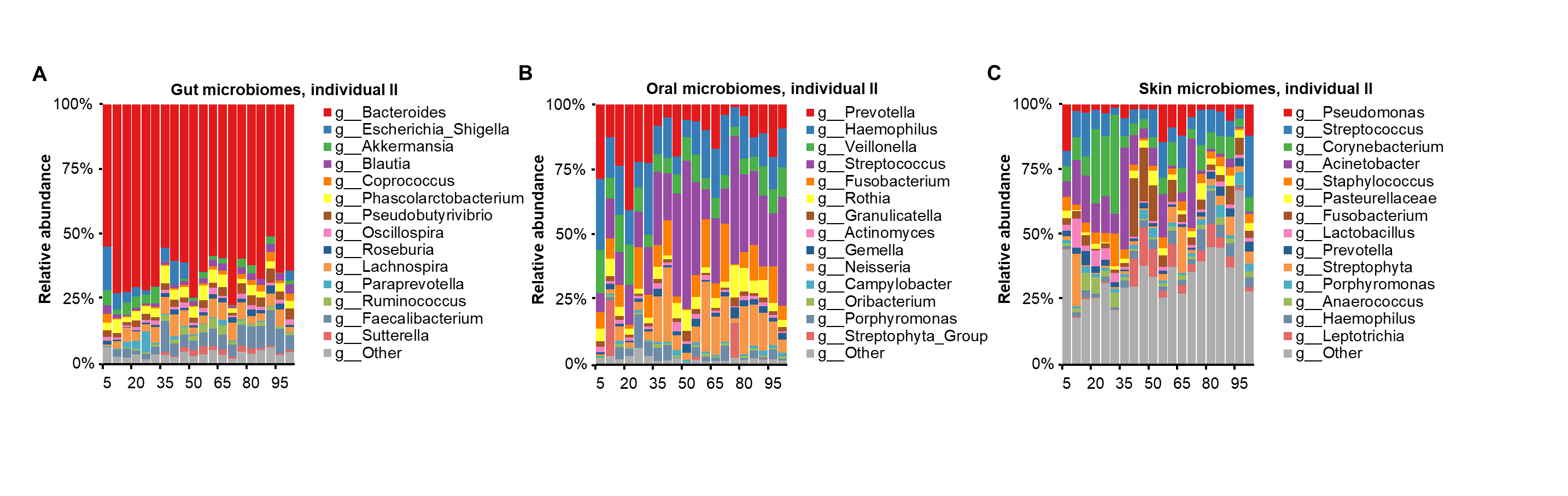

**Fig. S5. Transition of human microbiome across time and body sites.** The within-habitat microbiome transition of gut (**A**), oral (**B**) and skin (**C**) of individual II across 396 time points.

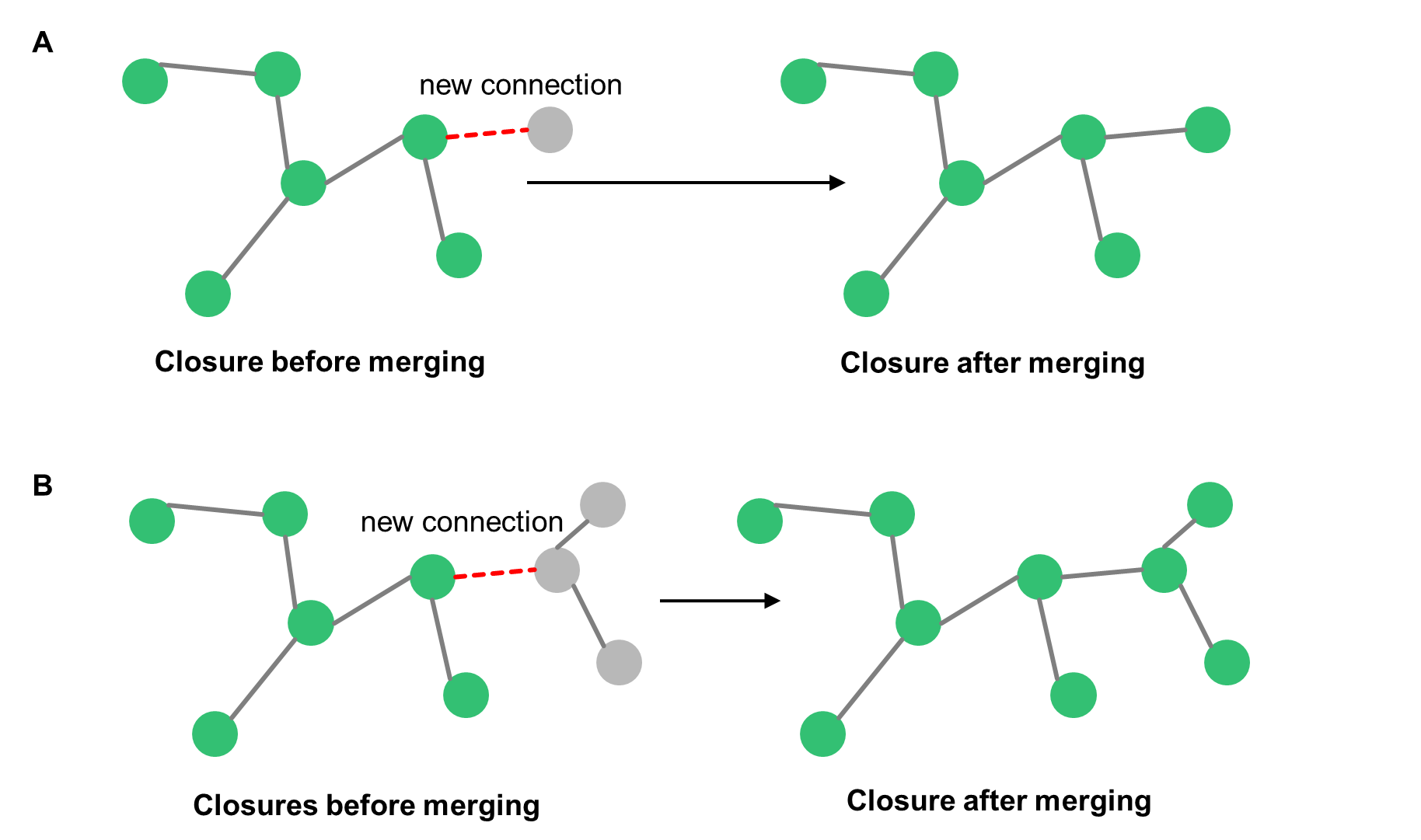

**Fig. S6. Expansion of a transitive closure by merging (A) linked node and (B) linked closure.**
